# Proximity labeling reveals a new *in vivo* network of interactors for the histone demethylase KDM5

**DOI:** 10.1101/2022.11.20.517232

**Authors:** Matanel Yheskel, Simone Sidoli, Julie Secombe

## Abstract

**Background:** KDM5 family proteins are multi-domain regulators of transcription that when dysregulated contribute to cancer and intellectual disability. KDM5 proteins can regulate transcription through their histone demethylase activity in addition to demethylase-independent gene regulatory functions that remain less characterized. To expand our understanding of the mechanisms that contribute to KDM5-mediated transcription regulation, we used TurboID proximity labeling to identify KDM5-interacting proteins.

**Results:** Using *Drosophila melanogaster*, we enriched for biotinylated proteins from KDM5-TurboID-expressing adult heads using a newly generated control for DNA-adjacent background in the form of dCas9:TurboID. Mass spectrometry analyses of biotinylated proteins identified both known and novel candidate KDM5 interactors, including members of the SWI/SNF and NURF chromatin remodeling complexes, the NSL complex, Mediator, and several insulator proteins.

**Conclusions:** Combined, our data shed new light on potential demethylase-independent activities of KDM5. In the context of KDM5 dysregulation, these interactions may play key roles in the alteration of evolutionarily conserved transcriptional programs implicated in human disorders.

## Background

Lysine demethylase 5 (KDM5) family proteins are multidomain transcriptional regulators able to recognize and enzymatically modify chromatin.(1,2) The best characterized function of KDM5 proteins is their histone demethylase activity, which cleaves a chromatin mark that is found at most active promoters, trimethylated lysine 4 on histone H3 (H3K4me3).(1,3–6) KDM5 proteins are evolutionarily conserved, with four paralogous genes in mammals encoding KDM5A-D, while animals with smaller genomes such as nematodes and flies possess a single *kdm5* gene. The importance of KDM5 function is emphasized by the observation that changes to the expression of this family of proteins is associated with two clinical outcomes: cancer and intellectual disability (ID).(7–9) KDM5A and KDM5B are amplified or overexpressed in a range of cancers, including breast, ovarian, skin, and lung.(8,10–13) KDM5A/B appear to play several roles in tumorigenesis, including promoting cell cycle progression and regulating the metabolism of cancer stem cells.(14–16) In contrast to the gain of function seen in cancer cells, loss of function variants in the autosomal paralogs KDM5A, KDM5B, and the X-linked KDM5C have been observed in individuals with intellectual disability.(17–21) KDM5 proteins have an evolutionarily conserved role in regulating critical gene expression programs in neurons as evidenced by morphological and functional neuronal phenotypes in KDM5B and KDM5C knockout mice.(21–23) Similarly, flies and nematodes with *kdm5* mutations display altered neuroanatomical development and neurotransmission. (24–26)

KDM5 catalytic function is mediated by the joint activity of the Jumonji N (JmjN) and JmjC domains and is classically thought to result in transcriptional repression. In addition, KDM5 proteins possess other potential gene regulatory domains, including plant homeodomain domain (PHD) motifs that can recognize H3K4me2/3 or H3K4me0, and a potential DNA binding A/T interaction domain (ARID).(27–32) These binding domains likely function in-concert with the histone demethylase activity of KDM5 by, for example, recruiting it to target promoters or altering enzymatic activity through the activity of individual or combinations of accessory domains. Conversely, non-enzymatic functions of these domains and/or other motifs of KDM5 that have no currently known function, such as the C_5_HC_2_ domain, could regulate transcription through distinct mechanisms. There is ample evidence that KDM5 proteins can regulate transcription independently of their demethylase activity. For instance, KDM5 is essential for viability in flies in a manner that is independent of its histone demethylase activity.(31,33) In addition, both demethylase-dependent and independent functions of KDM5 are critical for *Drosophila* neuronal development and function.(24,25) Consistent with this, some missense alleles of KDM5C observed in individuals with intellectual disability diminish its enzymatic activity, while others do not.(34–37) Similarly, demethylase dependent and independent activities of KDM5 proteins are likely to be important for their contributions to the etiology of and spread of cancers.(38,39) For example, KDM5B demethylase-independent functions in breast cancer promote metastatic potential to the lung.(38,39) Thus, even though KDM5 proteins derive their name from their enzymatic function, other conserved motifs contribute to their gene regulatory activities, although these activities remain much less characterized.

Understanding the repertoire of gene regulatory mechanisms utilized by KDM5 family proteins requires a comprehensive understanding of the proteins they can interact with. Traditional immunoprecipitation coupled with mass spectrometry (IP-MS) approaches have been used to identify proteins that form complexes with KDM5 family proteins in both mammals and *Drosophila*.(6,40–45) These experiments have revealed several conserved interactions, most notably with histone deacetylase 1 (HDAC1) and other proteins known to associate with this chromatin modifier.(42,44) To expand our understanding of the proteins that function with KDM5 to mediate its gene regulatory activities, we used TurboID-mediated proximity labeling.(46) This has been shown to be a powerful technique to identify weak or transient interactions that may otherwise be disturbed during the process of traditional IP experiments.(46–53) This technique takes advantage of the promiscuous biotin ligase activity of TurboID, which results in the biotinylation of lysine residues within 10 nm of its active site. When expressed as a chimeric fusion to a protein of interest, interacting proteins will be biotin-labeled.(46,54) Covalently modified proteins are then recovered with streptavidin beads and prepared for liquid chromatography-tandem mass spectrometry (LC-MS/MS). By expressing KDM5 that was N- or C-terminally tagged with TurboID *in vivo*, we recovered about half of previously identified interactions in *Drosophila*, and almost all interactions known to be conserved in mammalian cells, clearly demonstrating the robustness of this technique. Furthermore, we have discovered a novel interactome for KDM5 that suggests roles in the function of the switch/sucrose non-fermentable (SWI/SNF), non-specific lethal (NSL), nucleosome remodeling factor (NURF), and Mediator complexes, in addition to chromatin insulation.

## Results

### Chimeric TurboID-KDM5 proteins are functional and broadly biotinylate

To identify KDM5 interactors *in vivo*, we created constructs in which KDM5 was N- or C-terminally tagged with TurboID to maximally identify proteins that could function with KDM5. Because our long-term goal is to further develop our *Drosophila* model of KDM5-induced intellectual disability, we chose to carry out our TurboID studies using adult heads to enrich for neuronal tissue, using the general workflow shown in Fig 1A. Generating a TurboID system that closely mimics endogenous *kdm5* expression has been shown to be important for delivering more specific biotinylation compared to overexpression.(55) Based on our prior generation of a UASp-*kdm5* transgene that is expressed at approximately endogenous levels in somatic cells when crossed to a range of Gal4 drivers, we generated transgenic flies harboring HA-tagged UASp-*TurboID*:*kdm5* and UASp-*kdm5*:*TurboID* (Fig. 1B).(56) Therefore, we generated both N-terminal (NT-KDM5) and C-terminal (CT-KDM5) TurboID fusions of KDM5 to understand the full breadth of its interactome and to highlight terminus-specific interactions. To test the functionality and expression of the chimeric KDM5-TurboID proteins, we expressed them ubiquitously in a *kdm5*^*140*^ null mutant background.(33) Western blot analysis using adult heads showed that NT-KDM5 and CT-KDM5 were expressed at levels similar to those observed from an endogenously HA-tagged KDM5 (Fig. 1C). Importantly, N- and C-terminally tagged KDM5 proteins were able to restore viability to *kdm5*^*140*^ null mutant flies, which normally die prior to adulthood (Fig. 1D).(33) Tagging KDM5 with TurboID therefore does not interfere with its essential functions. Flies in which TurboID-KDM5 was the only source of KDM5 (*kdm5*^*140*^;Ubi-Gal4>*TurboID*:*kdm5*) were used for all subsequent experiments to maximize the number of interactors identified.

**Figure 1:**
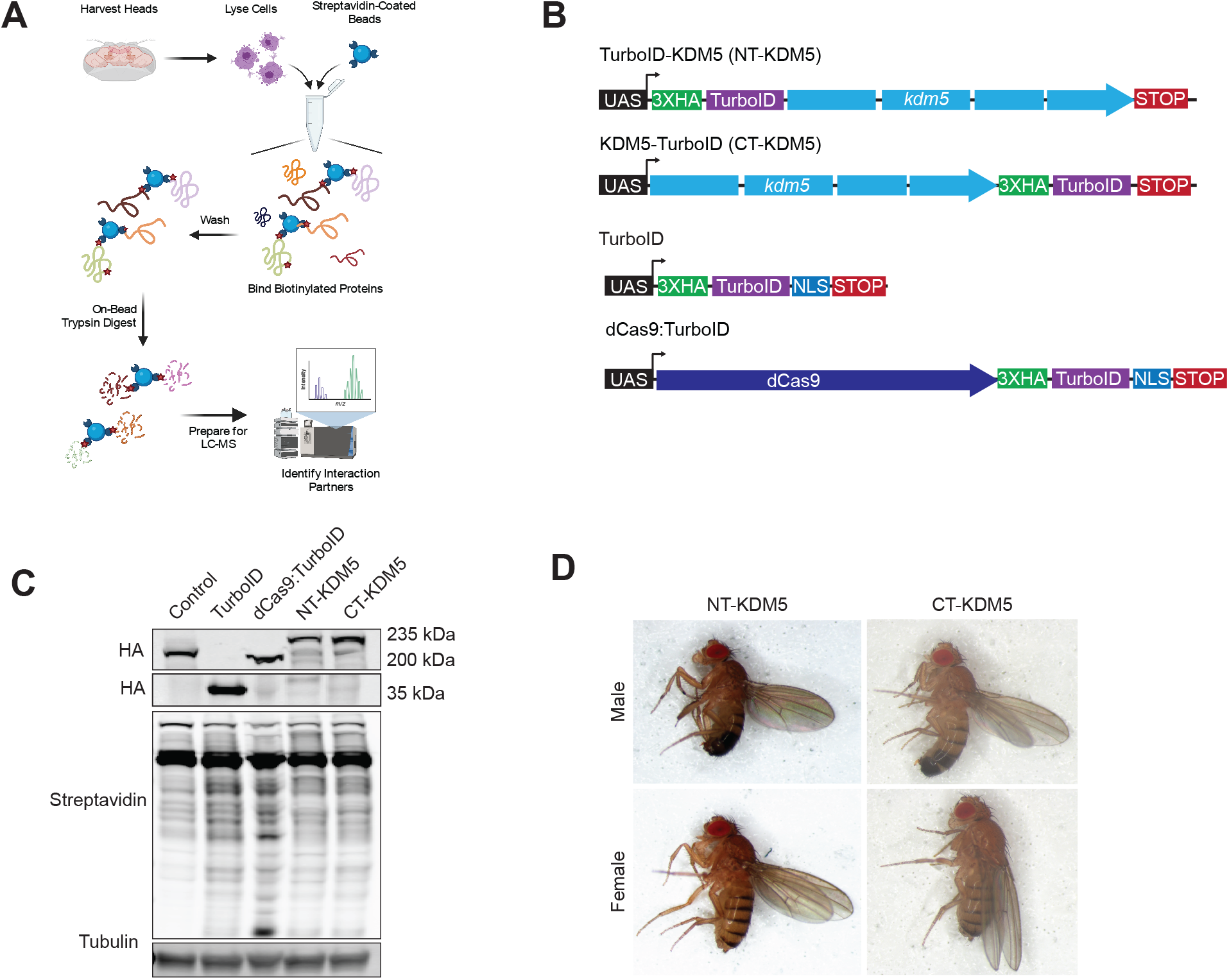
TurboID-tagged KDM5 proteins are functional and biotinylate endogenous proteins. (A) Schematic of the workflow used for purifying and fingerprinting biotinylated proteins from adult heads using LC-MS/MS mass spectrometry. (B) Schematic of the four UASp constructs generated able to express HA-tagged TurboID alone, dCas9:TurboID, NT-KDM5 and CT-KDM5. (C) Western blot using adult heads showing levels of expression of KDM5 using anti-HA, biotinylation using a streptavidin-680 conjugate and the loading control alpha-tubulin. Genotypes: *kdm5*^*140*^; *gkdm5:HA* (a wild-type strain; Control), Ubi-Gal4/+; UASp-*TurboID*/+ (TurboID), Ubi-Gal4/+; UASp-*dCas9:TurboID* (dCas9:TurboID), *kdm5*^*140*^, Ubi-Gal4/*kdm5*^*140*^; UASp-NT-*kdm5*/+ (NT-KDM5) and *kdm5*^*140*^, Ubi-Gal4/*kdm5*^*140*^; UASp-CT-*kdm5* /+ (CT-KDM5). (D) Rescue of *kdm5*^*140*^-induced lethality by ubiquitous expression of UAS-NT-*KDM5* or UAS-CT-*KDM5* using Ubi-Gal4. Genotype of male and female adult flies shown is *kdm5*^*140*^, Ubi-Gal4/*kdm5*^*140*^; UASp-NT-*kdm5* /+ (NT-KDM5) and *kdm5*^*140*^, Ubi-Gal4/*kdm5*^*140*^; UASp-CT-*kdm5* / + (CT-KDM5).

### Determining the proper controls to identify the KDM5 proximitome

To confidently identify proteins that function with KDM5, appropriate controls are critical. Due to the novelty of the technique, there are no standard controls for proximity labeling experiments. Many studies simply enrich over endogenous biotinylation and bead background.(57–60) Other studies expressed forms of TurboID alone that were localized to the specific cellular compartment that the protein of interest resided, such as the cellular membrane.(46,49,50) As a non-TurboID-expressing wild-type control with a similar genetic background, we used a fly strain in which endogenous *kdm5* is removed and HA-tagged KDM5 is expressed using its endogenous promoter from a transgene inserted at the same locus as the TurboID constructs (*kdm5*^*140*^;*gkdm5*^*WT*^). We will refer to this genotype as control. We also generated a transgene able to express nuclear localized, HA-tagged, TurboID alone using the same UAS promoter used for CT-*kdm5* and NT-*kdm5*, in an effort to assay general nuclear background (Fig. 1B). To compare the levels of TurboID alone to TurboID-KDM5 we expressed these transgenes using Ubi-Gal4 in a wild-type and *kdm5*^*140*^ background, respectively. Anti-HA western blot from adult heads showed significantly higher levels of expression for TurboID alone, possibly creating high levels of background biotinylation in this strain (Fig. 1C). Because biotin is essential for animal viability and thus included in the standard fly food used for crosses and stock maintenance, we assessed the ability of all TurboID transgenes to biotinylate proteins when expressed using Ubi-Gal4 by probing with infrared-conjugated streptavidin. Compared to control flies, similar levels of biotin-conjugated proteins were observed in heads expressing TurboID, NT-KDM5 and CT-KDM5, demonstrating their ability to biotinylate *in vivo* (Fig 1C).

To identify proteins preferentially biotinylated by NT-KDM5 and CT-KDM5 compared to control and TurboID alone, we carried out streptavidin-bead pulldowns in quadruplicate followed by LC-MS/MS. This experiment (experiment 1) identified a total of 1332 proteins, 476 of which are found in the nucleus where we have previously shown KDM5 to be localized.(25) Principal component analysis (PCA) of normalized nuclear protein abundances showed that TurboID alone, NT-KDM5, and CT-KDM5 clustered together, but were distinct from controls (Fig S1A). We therefore compared the proteins identified in NT-KDM5 and CT-KDM5 to control heads which revealed enrichment of 172 and 184 proteins, respectively, using a *p*-value cutoff of 0.05 (Fig S1B, C; Table S1). 136 proteins were commonly enriched by N- and C-terminally tagged KDM5, suggesting that we can robustly detect proteins in proximity to KDM5 (Fig. S1D). To assess the quality of our data, we determined how many known KDM5 interactors were identified in our analyses. Fifteen proteins have been established to form a complex with *Drosophila* KDM5 through IP-MS studies or targeted co-IP experiments (Table S2). Suggesting the robustness with which the TurboID approach identifies *bona fide* KDM5-associated proteins, 7 known interactors were identified by NT-KDM5 and 6 by CT-KDM5 (47% and 40%, respectively). We additionally assessed biotinylated protein enrichment of NT-KDM5 and CT-KDM5 compared to TurboID alone (Fig S1E, F). As expected, based on the increased level of expression of this protein compared to TurboID-tagged KDM5, a high level of background was observed in these flies. This resulted in fewer proteins being enriched in NT-KDM5 and CT-KDM5 (32 and 61, respectively), reduced overlap between the datasets, and a reduction in the number of known interactors identified (Fig S1G, H). Interestingly, TurboID alone appears to show bias in its biotinylation of nuclear proteins, as comparing TurboID to control revealed significant enrichment of 199 proteins, a majority of which are involved in chromatin-mediated transcriptional regulation (Fig. S1I).

Because of concerns related to the use of TurboID alone as a control, we generated a transgene encoding an enzymatically inactive form of the Cas9 enzyme (dCas9) fused to an HA-tagged nuclear localized TurboID (UASp-*dCas9*:*TurboID*). The encoded dCas9:TurboID fusion protein is more similar in size to the KDM5 fusion proteins, being 194kDa and 235kDa, respectively, compared to 36kDa for TurboID alone. In addition, dCas9 can scan the DNA, potentially making this fusion an appropriate control for chromatin binding proteins such as KDM5 by restricting biotinylation to DNA-adjacent proteins.(61) Ubi-Gal4-mediated expression of this transgene revealed that dCas9:TurboID was expressed at similar levels to the KDM5-TurboID fusion proteins and was able to biotinylate (Fig. 1C). Repeating the proximity labeling experiment, we carried out triplicate streptavidin-bead pulldowns from heads of control, TurboID, dCas9:TurboID, NT-KDM5 and CT-KDM5 flies (experiment 2). MS analyses identified 1,146 proteins across all samples, 203 of which were nuclear. PCA from this second experiment showed that NT-KDM5 and CT-KDM5 clustered together, indicating that these datasets are more alike to each other than to any of the controls (Fig S2A). Like our first experiment, TurboID alone clustered with NT-KDM5 and CT-KDM5, and was distinct from control and dCas9:TurboID samples. Using these data, we compared NT-KDM5 and CT-KDM5 to control, dCas9:TurboID, and to TurboID alone (Table S3). Proteins enriched in the KDM5 samples compared to control gave similar results to those obtained in the first experiment (Fig 2A, B). Using control flies as reference, 82 proteins were identified using NT-KDM5 and 68 for CT-KDM5, with 61 of these proteins being identified in both datasets. Compared to TurboID alone, only 29 and 24 proteins were enriched for NT-KDM5 and CT-KDM5, with 13 overlapping between the two datasets (Fig. 2C, D). Importantly, we find that comparing NT-KDM5 and CT-KDM5 with dCas9:TurboID yielded data very similar to that seen using control animals, despite providing a higher biotinylation background. 66 and 59 proteins were enriched in NT-KDM5 and CT-KDM5, respectively, with an overlap of 48 proteins (Fig. 2E, F).

**Figure 2:**
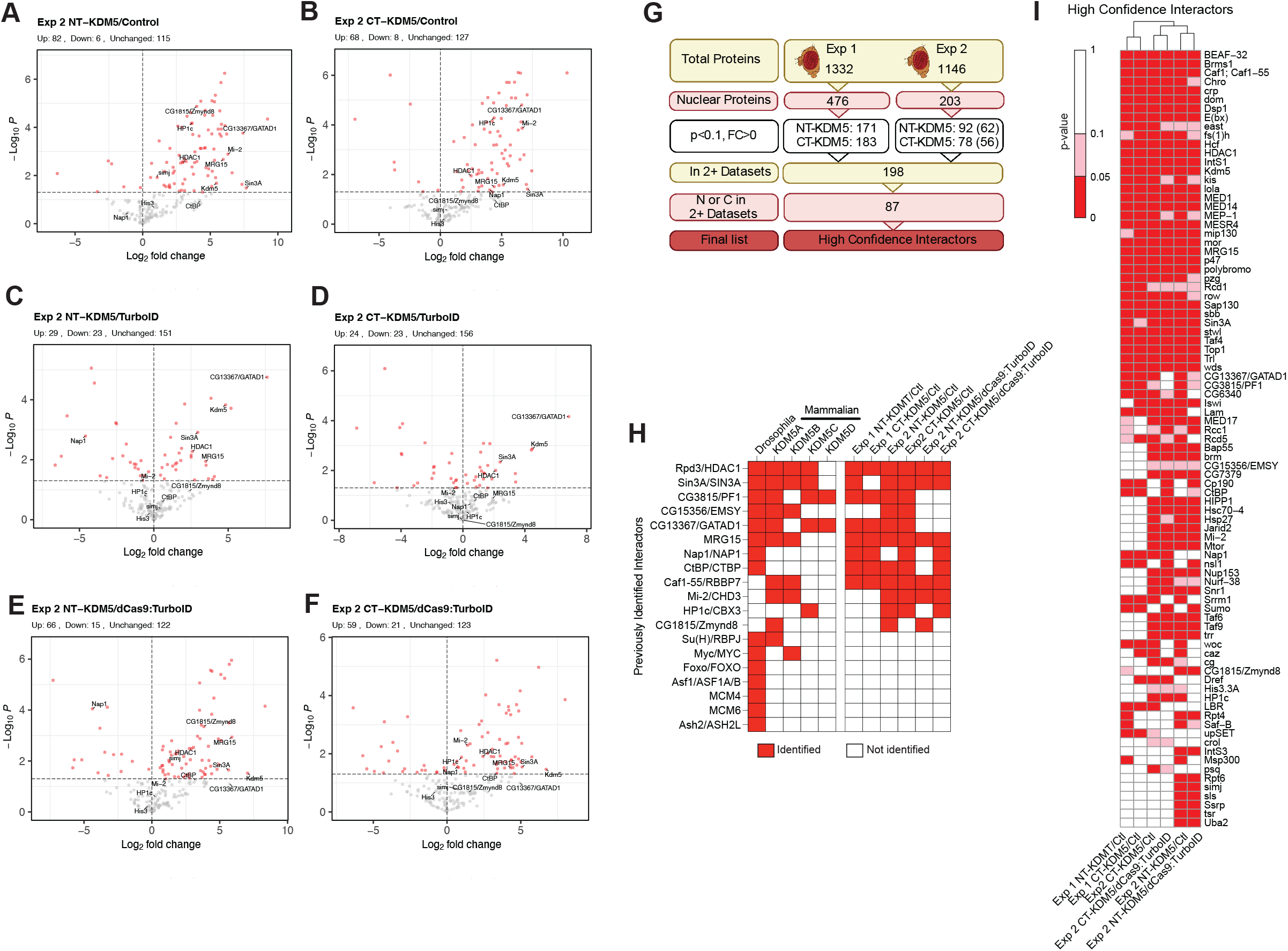
Identification of high confidence KDM5 interactors through TurboID. (A) Volcano plot showing data comparing biotinylated proteins enriched by NT-KDM5 to control. (B) Volcano plot showing data comparing CT-KDM5 to control. (C) Volcano plot showing data comparing NT-KDM5 to TurboID. (D) Volcano plot showing data comparing CT-KDM5 to TurboID. (E) Volcano plot showing data comparing NT-KDM5 to dCas9:TurboID. (F) Volcano plot showing data comparing CT-KDM5 to dCas9:TurboID. (G) Filtering workflow for identification of high confidence KDM5 interactors by combining data from experiments 1 and 2. (H) Summary of known *Drosophila* KDM5 interactors, whether the interaction is conserved in mammals (mouse or human) and the identification of these proteins in experiment 1 and experiment 2 compared to control and dCas9:TurboID. Three interactors identified in mammalian cells but not previously in *Drosophila* are also included (Caf1-55, Mi-2 and Zmynd8). (I) High confidence interactors (HCI) based on their identification in experiment 1 (compared to control) and experiment 2 (compared to control and dCas9:TurboID). Dark red box indicates enrichment of *p*<0.05, pink indicates *p*<0.1. For all volcano plots shown, known interactors are indicated with text. Red dots indicate significantly enriched proteins (*p*<0.05) and the dotted line on Y-axis indicates *p*=0.05.

To complete our characterization of dCas9:TurboID as a tool to identify enriched DNA-adjacent proteins, we compared these data to both TurboID alone and to control. Similar to data from the first experiment comparing TurboID and control, TurboID alone shows a preference for biotinylating a large number of chromatin-related proteins, even compared to dCas9:TurboID (Fig. S2B, C). Comparing dCas9:TurboID to control revealed enrichment in a relatively small number of proteins that were enriched for transcriptional-regulatory proteins consistent with the ability of dCas9:TurboID to biotinylate targets while scanning DNA (Fig. S2D). Confirming the challenges related to using TurboID alone, the consistency with which proteins were enriched in NT-KDM5 or CT-KDM5 compared to TurboID alone was very low, with little agreement across experiments using a *p*-value cutoff <0.05 or <0.1 (Fig. S2E, F). Moreover, the number of previously identified interactors remained low in experiment 2 when comparing to TurboID alone, with only 5 and 3 being identified in NT-KDM5 and CT-KDM5, respectively (Fig. S2G). We therefore suggest that endogenous biotinylation or dCas9:TurboID are superior to TurboID as controls for proximity labeling experiments where the protein of interest is nuclear-specific and DNA-adjacent.

### Proximity labeling identifies new potential KDM5 interacting complexes

To build a high confidence list of proteins which interact with KDM5, we combined data from experiment 1 comparing NT-KDM5 and CT-KDM5 to control (2 datasets), and experiment 2 in which NT-KDM5 and CT-KDM5 were compared to control (2 datasets) as well as dCas9:TurboID (2 datasets). To do this, we began by filtering for enriched nuclear proteins across all six datasets using a *p*-value<0.1. We first filtered for proteins identified in at least two of six datasets. Then to include the possibility of terminus-exclusive interactors, we required that at each terminus proteins had to be identified in 0 (exclusive to other terminus) or in 2 of 3 datasets (Fig. 2G). Demonstrating the power of this approach, this included 7 of the 15 (47%) known *Drosophila* KDM5 interactors (Fig. 2H, I). This ratio increased to 7 of 8 (88%) for those interactors that have been shown in both *Drosophila* and mammalian cells. In addition, we identified several proteins not previously been found to be *Drosophila* KDM5 interactors but have been purified with mammalian KDM5A, KDM5B, KDM5C and/or KDM5D. These included the nucleosome remodeler Mi-2, the chromatin assembly factor 1 (Caf1;Caf1-55), the actyl-lysine binding protein Zmynd8 and the heterochromatin-associated protein HP1c (Fig 2H, I). (6,42,44) With these stringent filtering criteria, we identified a total of 87 proteins (Fig. 2I).

Our proximity-labeling studies using Turbo-KDM5 revealed a broader interactome than previously described in the literature. To better understand the relationships between the proteins identified in our study, we generated a protein interaction map using Cytoscape and STRING (Fig. 3A).(62,63) Gene Ontology (GO) analysis shows that many of these proteins have roles in the regulation of gene expression, chromatin modification, and chromatin remodeling (Fig. 3B). In addition to confirming the strong link between KDM5 and Sin3/HDAC1-containing complexes, these analyses also highlighted interactions with new protein complexes. Among these, we find proteins such as Boundary Element-Associated Factor of 32kDa (BEAF-32), Chromator (Chro), Putzig (Pzg), and Centrosomal protein 190kDa (Cp190) that function in regulating genomic architecture, suggesting a unstudied role for KDM5 in this process.(64–69) In addition, we identified proteins critical for forming the transcriptional pre-initiation complex (TPIC), which is consistent with the promoter-proximal binding of KDM5 proteins across species.(25,44,70–72) Using a recent cryo-EM structure of the human TPIC, we found that distinct surfaces interacted with KDM5, consistent with the specificity of biotinylation using TurboID-KDM5.(73) Specifically, three adjacent proteins in the mediator complex (MED1, MED14 and MED17) and three adjacent subunits of TFIID (Taf4, Taf6, and Taf9) were identified in our analyses (Fig. 3C). This suggests that KDM5 may play a role in enhancer-promoter communications that regulate the transcriptional activity of target genes.

**Figure 3:**
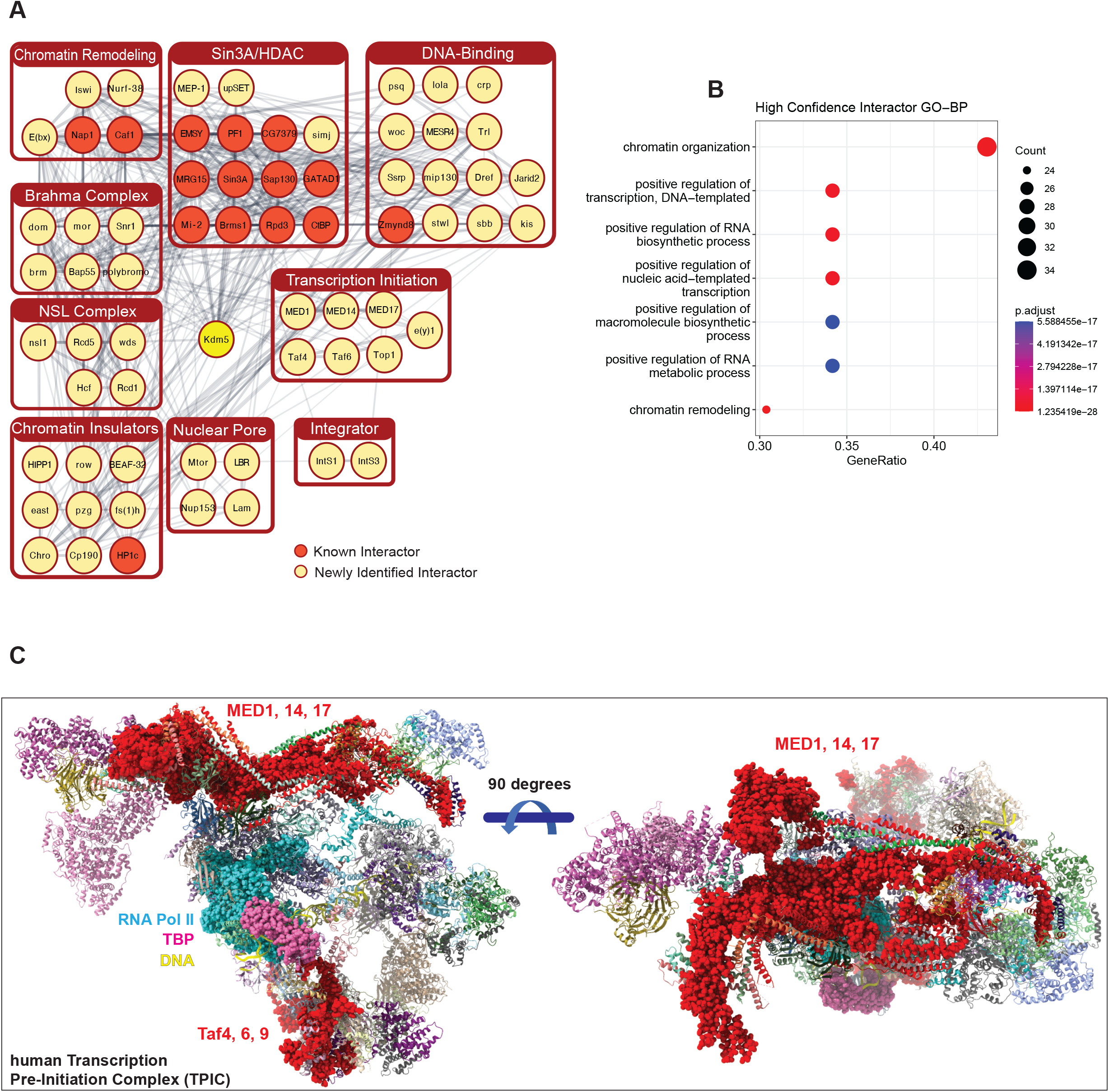
TurboID-tagged KDM5 biotinylates proteins involved in several aspects of gene expression regulation. (A) STRING analyses of nuclear proteins that were significantly biotinylated by NT-KDM5 and CT-KDM5. Grey lines indicate known physical interactions between proteins. Darker lines indicate a higher confidence of interaction. Cytoscape was used to manually cluster annotated proteins based on their STRING Cluster Enrichment and known functions based on published literature. Unconnected nodes and proteins with unclear links to known complexes are not shown. (B) Gene Ontology Biological Process (GO-BP) analyses of the 87 high confidence KDM5 interacting proteins. (C) Structure of the human pre-initiation complex (PDB accession: 7ENA) showing proteins identified as high confidence interactors in our TurboID data in red bubbles. DNA (yellow), TBP (pink bubbles) and RNA polymerase II (cyan bubbles) are also shown.

### KDM5 and newly identified interactors occupy overlapping genomic binding sites in

#### *Drosophila* and human cells

To further explore the relationship between KDM5 and newly identified interactors in *Drosophila*, we compared their genomic binding with those of interactors using publicly available ChIP-seq datasets. We used published KDM5 ChIP-seq data from whole adult flies to interrogate ChIP-Atlas as a means to identify datasets from any *Drosophila* cell type that significantly overlap (via permutation 100X).(74) We then overlapped these with our high confidence interactor (HCI) list to reveal a total of 27 overlapping datasets. 30 proteins in our interactor list had available data on ChIP Atlas. (Fig. 4A, B). These overlapping datasets included known interactors such as Sin3A, in addition to new interactors BEAF-32, the DNA replication-related element factor Dref and the NURF chromatin remodeler component Iswi. We analyzed the distribution of KDM5 around interactor peaks and found that KDM5 seems to flank their binding sites (Fig. 4C-F). In these cases, the distribution of KDM5 appears to be bimodal while Sin3A, BEAF-32, Dref and Iswi have a single peak. This is likely due to KDM5 binding to promoter regions of adjacent genes with divergent promoters, leading to two peaks occurring within the 4kb range shown.(75) Sin3A, BEAF-32, Dref, and Iswi bind to a single site that overlaps with the region bound by KDM5 at one or both promoters. In contrast, KDM5 and female sterile (1) homeotic (fs(1)h), which encodes the ortholog of the acetyl-histone binding Brd2/Brd4, appear coincident (Fig. 4G). It is also notable that for Sin3A and Iswi, KDM5 does not co-localize across all binding sites (Fig. 4C, F). This could simply reflect binding differences in the cell types used for the ChIP-seq studies, or that specific promoter sub-types are co-occupied. A combined genome browser snapshot highlights the binding of KDM5, Sin3A, BEAF-32, Dref, Iswi and fs(1)h relative to each other, and also relative to the transcriptional start site (TSS; Fig. 4H; Fig. S3A).

**Figure 4:**
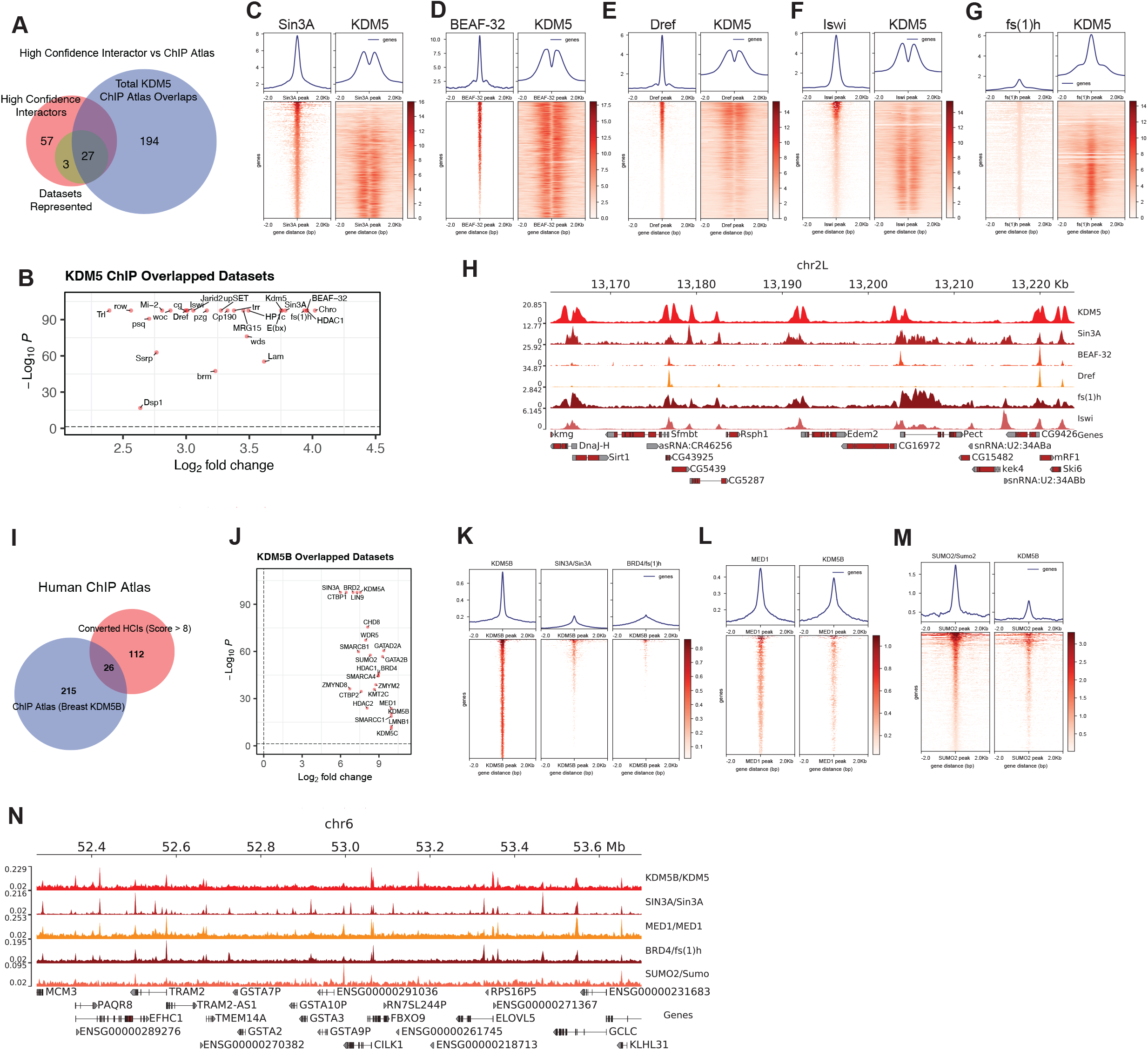
Proteins enriched in KDM5 TurboID experiments show overlapping genomic binding profiles in *Drosophila* and human cells. (A) Venn diagram showing overlap between high confidence biotinylated proteins (red) and datasets enriched when comparing published KDM5 ChIP-seq data from whole adult flies to other *Drosophila* ChIP datasets (blue). A total of 27 datasets showed significant overlap using ChIP-atlas. 30 interactors had ChIP-seq datasets in the ChIP Atlas database (green). (B) Volcano plot showing fold enrichment and p-values of the permutation analyses between KDM5 and 27 ChIP-seq datasets of high confidence interactors. (C) Heat maps showing ChIP-seq genomic binding profiles of Sin3A from S2 cells and KDM5 from whole adult flies. (D) Genomic binding profiles of BEAF-32 ChIP-seq from S2 cells and KDM5. (E) Genomic binding profiles of Dref ChIP-seq from Kc167 cells and KDM5. (F) Genomic binding profiles of Iswi ChIP-seq from Kc167 cells and KDM5. (G) Genomic binding profiles of fs(1)h ChIP-seq from Kc167 cells and KDM5. (H) Representative genome browser image showing binding of KDM5, Sin3A, BEAF-32, Dref, fs(1)h, and Iswi. (I) Venn diagram showing overlap between human orthologs of *Drosophila* KDM5 high confidence interactors (HCI) and their enrichment when comparing published KDM5B ChIP-seq data to breast cancer ChIP-seq datasets using ChIP-Atlas. 26 datasets significantly overlapped. (J) Volcano plot showing fold enrichment and p-values of the permutation analyses between KDM5B and 26 ChIP-seq datasets of human-ortholog converted high confidence interactors. (K) Genomic binding profiles of KDM5B, SIN3A and BRD4 showing similar genome-wide binding. Binding is shown relative to KDM5B due to the much larger number of SIN3A and BRD4 binding sites in the genome compared to KDM5B. (L) Genomic binding profiles of MED1 ChIP from MCF-7 cells and KDM5B showing similar localization. (M) Genomic binding profiles of SUMO2 (ortholog of *Drosophila* Sumo) ChIP from MCF-7 cells and KDM5B showing similar localization. (N) Representative genome browser image showing binding of KDM5B, SIN3A, MED1, BRD4 and SUMO2.

To assess the extent to which our high confidence KDM5 interactors might be evolutionarily conserved, we investigated genomic co-occupancy in human cells. We first converted our 87 *Drosophila* high confidence interactors to their human ortholog(s), which resulted in a total of 138 proteins due to humans possessing multiple paralogous proteins for some *Drosophila* proteins (DIOPT v8.0 score > 8/15; Table S4).(76) Due to the strong link between KDM5B and breast cancer, and the wealth of ChIP-seq datasets available in cell lines derived from this cancer type, we used KDM5B data from MCF-7 cells for these analyses.(77) Using peaks called from this ChIP-seq data, we interrogated all available breast cancer cell line datasets again using ChIP-Atlas (100X permutation). This revealed that 26 candidate interactors had binding profiles that significantly overlapped with KDM5B binding (Fig. 4I, J). Interestingly, these included the KDM5 paralogs KDM5A and KDM5C, suggesting that there may be overlapping or redundant function for these proteins. Similar to our studies using *Drosophila*, some proteins identified were known interactors, such as SIN3A that is known show similar genomic binding to KDM5B (Fig. 4K).(78) Other proteins overlapped with our genomic binding studies in *Drosophila*, including Brd4 (fs(1)h), while other proteins were identified because datasets were available in human cells and not *Drosophila*. These included the Mediator subunit MED1 and the small ubiquitin-like modifier SUMO2 (*Drosophila* Sumo) (Fig. 4L, M). A combined genome browser emphasizes the colocalization of these proteins with KDM5B and their relationship to the TSS (Fig. 4N; Fig. S3B). These data also highlight the difference in genome size between *Drosophila* and human cells, with the greater distance between promoters resulting in a single binding peak. Combined, our data show that the high confidence KDM5 interactors identified in *Drosophila* may be important for the function of KDM5B and other KDM5 paralogs in mammalian cells.

### Identified KDM5 interactors are implicated in neurodevelopmental disorders

Based on the association between genetic variants in KDM5A, KDM5B, and KDM5C and ID and autism spectrum disorder (ASD), KDM5-interactors could potentially be implicated in neurological disorders.(22,26,79–82) To examine this in more detail, we used the Simons Foundation Autism Research Initiative (SFARI) genes as an up-to-date source of genes with significant causal links to ASD.(83) The 1231 genes in this database are scored based on the level of confidence of association, with a score of 1 being the strongest link, in addition to whether ASD occurs as part of a syndrome (S). Using our list of 138 human ortholog-converted KDM5 interactors, we found that 26 of these overlap with SFARI ASD-associated proteins (*p*=8e-08; Fig. 5A-B). Some of these proteins have clear links to each other, such as TAF4 and TAF6 that are components of TFIID, while others are associated with numerous other aspects of transcriptional regulation. To look more broadly into the link between KDM5 interactors and neurodevelopmental disorders, we used the Developmental Brain Disorder Gene Database (DBD) which is a curated list of genes implicated in disorders such as ID, ASD, attention deficit hyperactivity disorder (ADHD) and schizophrenia.(84) 24 human-converted orthologs represented in DBD have been shown to contribute to ID, ASD, ADHD, and Schizophrenia (Fig. 5C). Unsurprisingly given the frequency that ID and ASD co-occur, 14 proteins were identified in both datasets, including KDM5B, KDM5C and BRD4. In addition, 9 ID-associated proteins were identified, including MED17, the NURD chromatin remodeling complex component GATAD2A and the C-terminal binding protein (CtBP) transcriptional repressor. Combined, these analyses expand our understanding of the potential network of proteins that function with KDM5 and provide new avenues for investigating the links between KDM5 family proteins and the etiology of neurodevelopmental disorders. A summary of our KDM5 interaction data highlighting proteins with known roles in transcriptional regulation is shown in Figure 5D.

**Figure 5:**
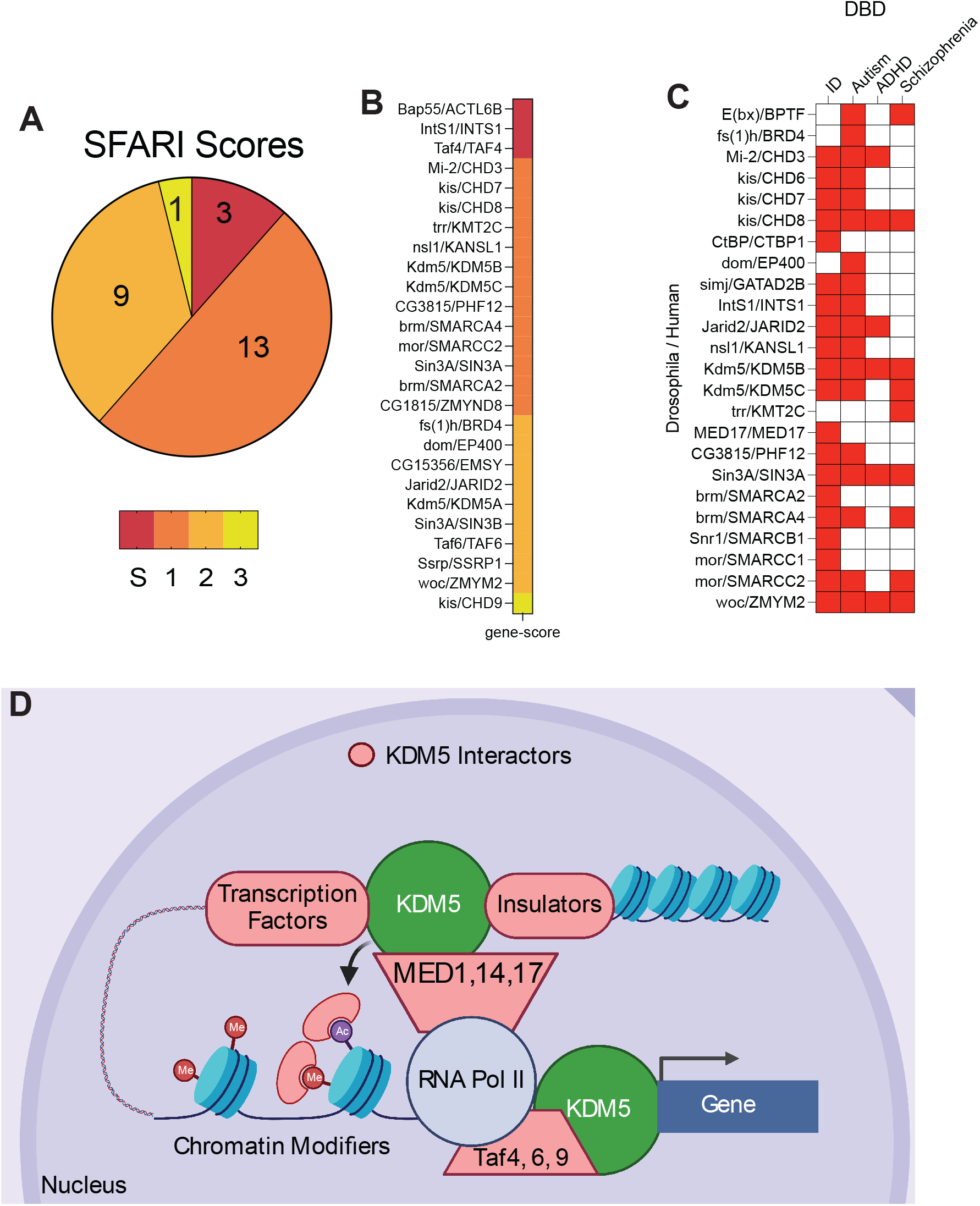
A subset of KDM5 interactors are implicated in neurodevelopmental disorders. (A) Overlap between human orthologs of *Drosophila* KDM5 interactors and genes associated with ASD using the SFARI database. S indicates syndromic ASD. Scores indicate confidence of causal association, with 1 indicating strongest link. (B) 26 candidate interacting proteins identified and their SFARI score. (C) 24 candidate interacting proteins were identified as being implicated in ID, ASD and/or schizophrenia using DBD. (D) Model for KDM5 interactions with key transcriptional proteins that are likely to impact the expression of downstream target genes.

## Discussion

Here we describe the interactome of *Drosophila* KDM5 in the adult head using TurboID-mediated proximity labeling. To identify the broadest selection of potential interactors, we N- and C-terminally TurboID-tagged KDM5 as some interactors were expected to be in proximity to both termini while others might be terminus specific. Importantly, TurboID-KDM5 chimeric proteins were functional, as they were able to rescue the lethality caused by a *kdm5*^*140*^ null allele. Using NT-KDM5 and CT-KDM5, we performed two experiments to optimize the experimental controls, as none are established for this relatively new technique. Our study revealed that expression of TurboID alone led to high background levels of biotinylation, particularly of chromatin-related proteins. In contrast, using control (endogenous biotinylation and bead background) or dCas9:TurboID provided similar and more reasonable background to identify proteins enriched by expression of TurboID-KDM5. Because we carried out two separate MS experiments, we were able to stringently filter our data to retain only those proteins that were nuclear localized and showed high reproducibility and rigor. This led to the identification of 87 high confidence KDM5 interactors, 12 of which were previously described in either *Drosophila* or mammals. Notably, while we refer to proteins identified in our TurboID analyses as interactors, we acknowledge that proximity labeling does not necessarily detect direct interactions. However, TurboID biotinylates lysine residues within 10 nm of its active site(54), which is equivalent to about 27 bp of B-DNA (8 bp per 3.4 nm). However unlikely, this short biotinylation radius can result in the identification of proteins that are nearby but in a distinct complex that do not physically touch KDM5. While these proteins are not in the same complex(es) as KDM5, they could still function with KDM5 to regulate gene expression by acting in concert with, or independently of, its histone demethylase activity. For simplicity, we will refer TurboID-enriched proteins as interactors.

Several lines of evidence allow us to have confidence in our described KDM5 interactome. The first is that we identified 88% of proteins previously described as KDM5 interacting proteins in flies and mammalian cells. We did not, however, detect all previously known KDM5 interactors in our proximity-labeling studies. For example, our prior studies have shown interactions between KDM5 and the transcription factors Myc and Foxo, and one of the best-established interactions of mammalian KDM5 proteins is with the Retinoblastoma protein (RBF in *Drosophila*).(5,72,85) None of these proteins were significantly enriched in our current study. Because these interactions have not been examined in neuronal cells, this may simply reflect differences in KDM5 complex composition across cell types. Alternatively, these complexes may be low abundance and therefore more difficult to detect by proteomic approaches. While we undoubtably missed some KDM5 interactors, we were able to reproducibly enrich a number of proteins using both NT- and CT-KDM5 across two independent experiments. In addition, many of the proteins identified have known physical connectivity with each other. Thus, rather than identifying individual components of complexes, we identified proteins well known to complex with each other, such as the SWI/SNF and NURF chromatin remodeling complexes. Interestingly, a functional link between KDM5 and these complexes is supported by studies in mouse embryonic stem cells which showed that a loss of KDM5B altered nucleosome position surrounding the TSS, although the mechanism was not revealed.(86) We also identified the insulator proteins BEAF-32, Chromator, Putzig, and Cp190 which complex together.(66,67) Our previous investigation in a fly strain harboring an allele associated with human intellectual disability identified enrichment of BEAF-32 binding sites at dysregulated genes.(26) Functionally, it is also notable that mutations in *kdm5, BEAF-32*, and *putzig* all modify position effect variegation (PEV) suggesting the possibility that these proteins function together to regulate chromatin compaction and/or organization.(87–89) For some TurboID-identified proteins such as the Mediator complex components (MED1, MED14, and MED17), as well as the TFIID proteins (Taf4, Taf6, and Taf9), published structural data are consistent with their link to KDM5.(73,90) The Mediator and Taf proteins identified neighbor each other, respectively, in the hTPIC cryo-electron microscopy structure. Moreover, we found that KDM5 interactions could be mapped to distinct surfaces at the hTPIC, suggesting one way that KDM5 could localize with respect to key transcriptional initiation machinery. Our enrichment of a subset of transcriptional preinitiation proteins implies that this is not simply due to KDM5 proteins binding near the promoter region of its target genes. If that were the case, then the entire preinitiation complex would have been identified in our datasets, including TBP and RNA Pol II. We additionally observed enrichment for proteins implicated in enhancer function, such as Zmynd8 that has previously been shown to interact with KDM5A and KDM5D and binds to monomethylated histone H3 lysine 4 (H3K4me1), a chromatin mark that is found at enhancers.(42,44) Consistent with the possibility that KDM5 may impact the chromatin status and activity of enhancers, our studies additionally revealed enrichment for the methyltransferase responsible for depositing H3K4me1, Trr/KMT2C.(91) Further studies are now required to define precisely which proteins directly interact with KDM5 to provide insight into how KDM5 carries out its functions to influence gene expression. Importantly, given the range of proteins found in our study, KDM5 may use distinct mechanisms to modulate gene expression levels in different genomic contexts and in cell distinct types. Although limited by the number of available ChIP-seq datasets available, corroborating evidence for our interactors also comes from the extensive overlap in genomic binding observed in *Drosophila* and/or mammalian cells.

The relationships between KDM5 and other gene regulatory complexes provide insight into how its dysregulation could contribute to human disorders. Many of the interacting complexes identified in our study have been implicated in tumorigenesis, including NSL and SWI/SNF.(92– 95) Furthermore, like KDM5, MED1 has been implicated as a transcriptional coactivator that mediates breast cancer metastasis and treatment resistance.(96,97) Identification of KDM5 interactors may provide insight to mechanisms of KDM5-mediated transcriptional regulation which underlie tumor development and progression. Changes to protein interactions could also contribute to the intellectual disability seen in individuals with genetic variants in KDM5A, KDM5B, or KDM5C. Indeed, for variants that do not alter histone demethylase activity, this may be a contributor to cognitive dysfunction. Our analyses of KDM5 interactors revealed an enrichment in proteins found to be altered in neurodevelopmental disorders whose clinical presentations overlap with those seen for KDM5 genes. KDM5 and interacting proteins could therefore influence neurodevelopment through common pathways. Altogether, our study suggests that KDM5 likely functions through numerous transient interactions with interconnected complexes to regulate gene expression in a context-dependent manner.

## AUTHOR CONTRIBUTIONS

Conceptualization, M.Y., J.S.; Methodology, M.Y., S.S.; Investigation, M.Y. and J.S.; Writing – original draft, J.S. and M.Y., Writing – Reviewing and Editing, J.S., M.Y., and S.S..; Funding acquisition, J.S., M.Y and S.S., Supervision, J.S. and S.S.

## DECLARATIONS

## Ethics approval and consent to participate

N/A

## Availability of data and materials

Transgenic fly strains described here are available upon request to Julie Secombe.

### ACKNOWLEDGMENTS

We thank members of the Secombe Lab, Melissa Castiglione, Blair Schneider, Michael Rogers, and Bethany Terry, as well as Michelle Schumacher for their feedback throughout this project and edits on the manuscript. We also appreciate the availability of stocks from the Bloomington *Drosophila* Stock Center (NIH P40OD018537) and are grateful to the Cancer Center Support Grant P30CA013330. This research was supported by the NIH Ruth L. Kirschstein National Research Service Award F31GM146347 and the Einstein MSTP Training Grant T32GM007288 to M.Y., NIH R01GM112783 and support from the Irma T. Hirschl Trust to J.S. The Sidoli lab gratefully acknowledges for financial support AFAR (Sagol Network GerOmics award), Deerfield (Xseed award), Relay Therapeutics, Merck, the NIH Office of the Director (1-S10-OD030286-01), the Einstein-Mount Sinai Diabetes Research Center, and the Einstein Cancer Center (P30-CA013330-47).

## FIGURE LEGENDS

**Figure S1:**
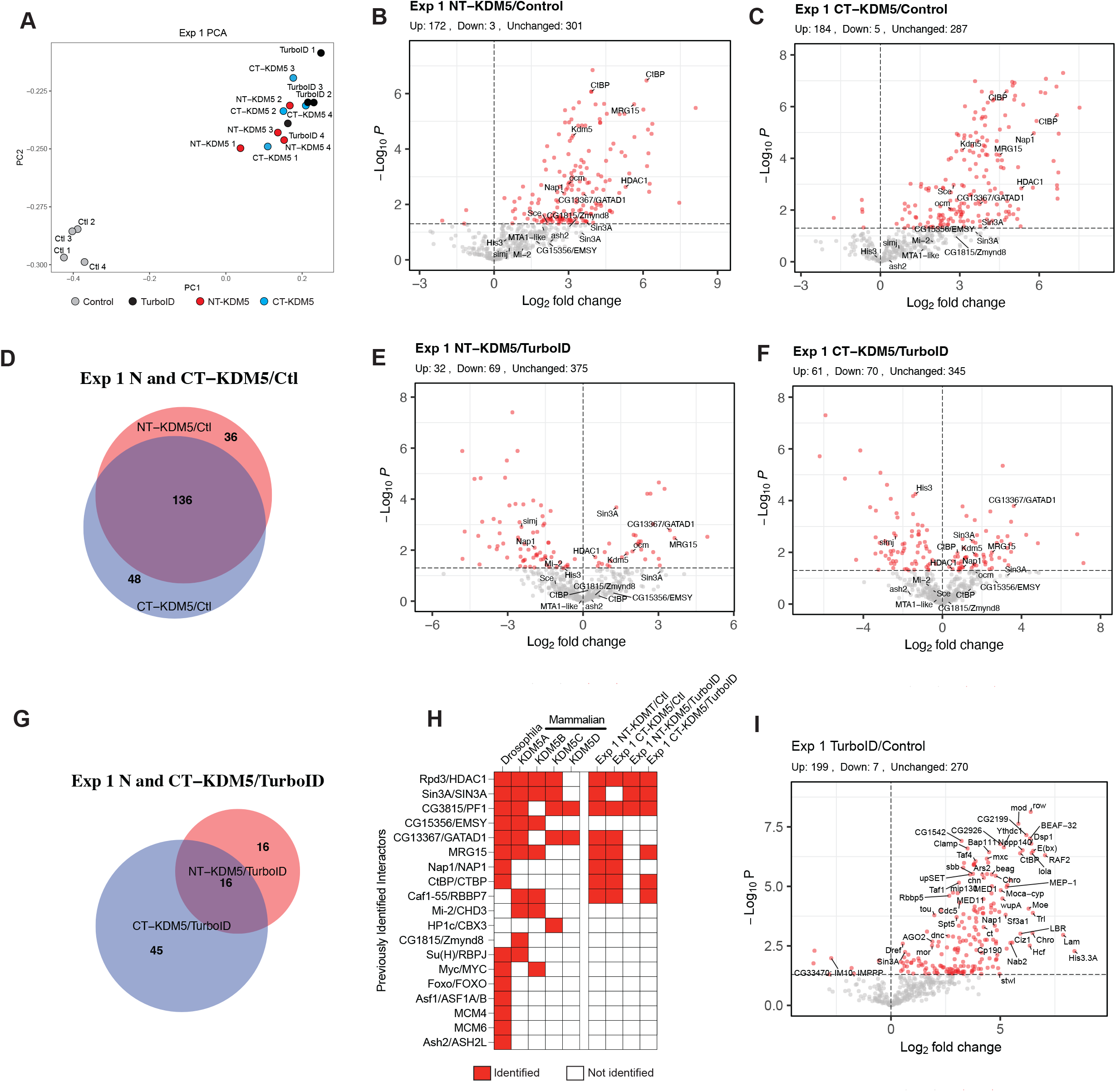
KDM5-TurboID studies using control and TurboID alone. (A) Principal component analysis of normalized nuclear protein abundances from control (Ctl), TurboID, NT-KDM5 and CT-KDM5 (experiment 1 data). (B) Volcano plot showing data comparing NT-KDM5 to control. (C) Volcano plot showing data comparing CT-KDM5 to control. (D) Venn diagram showing overlap of the proteins identified using NT-KDM5 and CT-KDM5 compared to control (Ctl). (E) Volcano plot showing data comparing NT-KDM5 to TurboID. (F) Volcano plot showing data comparing CT-KDM5 to TurboID. (G) Venn diagram showing overlap of the proteins identified using NT-KDM5 and CT-KDM5 compared to TurboID. (H) Summary of known *Drosophila* KDM5 interactors, whether the interaction is conserved in mammals (mouse and/or human) and the identification of these proteins in experiment 1 with all comparisons. Three interactors identified in mammalian cells but not previously in *Drosophila* are also included (Caf1-55, Mi-2 and Zmynd8). (I) Volcano plot showing data comparing TurboID to control. For all volcano plots shown, known interactors are indicated with text. Red dots indicate significantly enriched proteins (*p*<0.05) and the dotted line on Y-axis indicates *p*=0.05.

**Figure S2:**
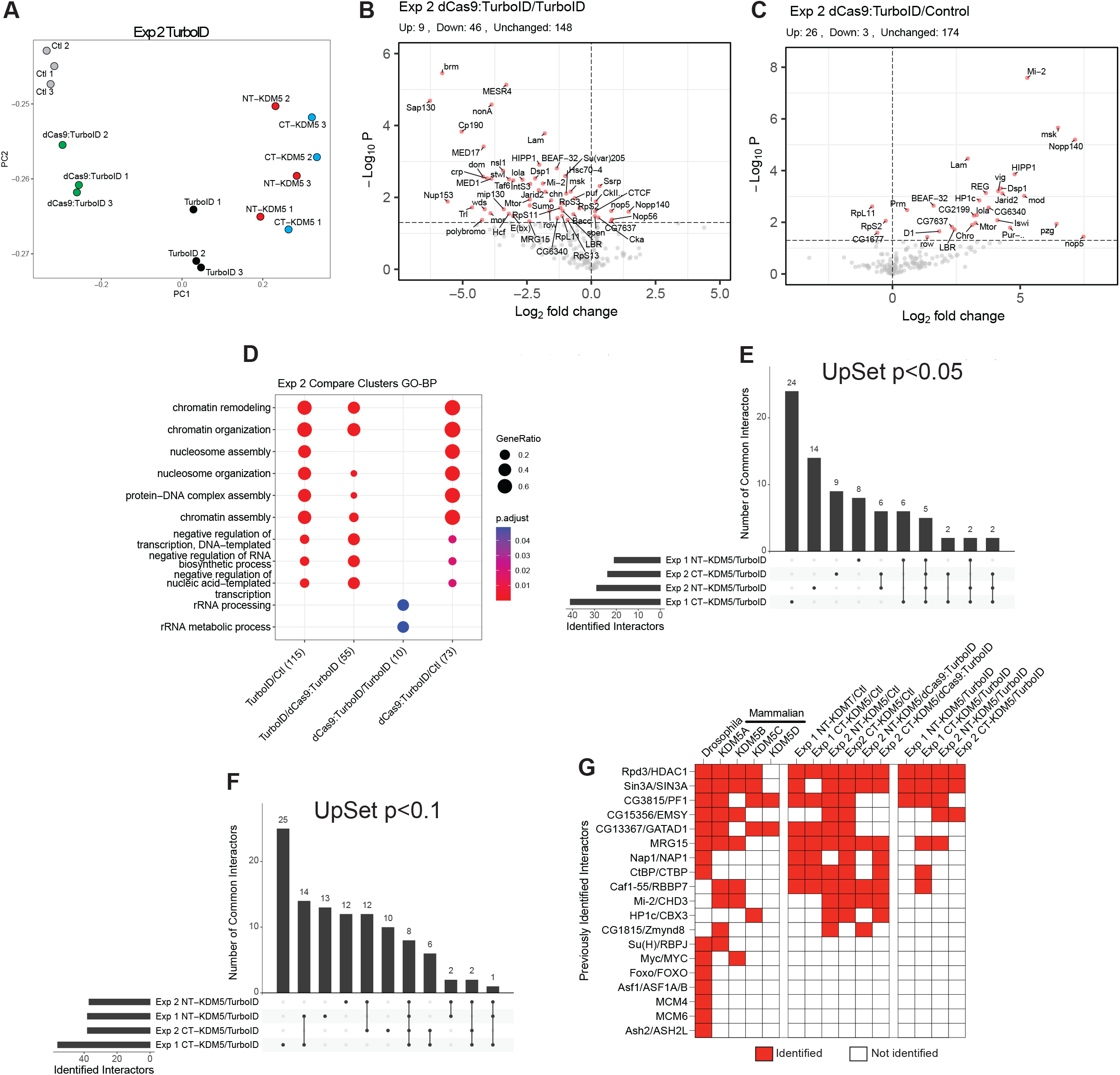
Comparing TurboID alone to dCas9:TurboID. (A) Principal component analysis of normalized nuclear protein abundances from control (Ctl; grey), dCas9:TurboID (T-dCas9; green), TurboID (black), NT-KDM5 (red) and CT-KDM5 (blue). (B) Volcano plot showing data comparing dCas9:TurboID to TurboID alone. Red dots indicate significantly enriched proteins (*p*<0.05) and the dotted line on Y-axis indicates *p*=0.05. (C) Volcano plot showing data comparing dCas9:TurboID to control. Red dots indicate significantly enriched proteins (*p*<0.05) and the dotted line on Y-axis indicates *p*=0.05. (D) GO analyses of proteins enriched comparing TurboID to control (Ctl), TurboID to dCas9:TurboID, dCas9:TurboID to TurboID and dCas9:TurboID to control. (E) UpSet plot showing the number of common interactors identified in experiments in which TurboID was used to compare datasets using *p*<0.05. (F) UpSet plot showing the number of common interactors identified in experiments in which TurboID was used to compare datasets using *p*<0.1. (G) Summary of all known interactors identified in all experimental comparisons from experiment 1 and experiment 2.

**Figure S3:**
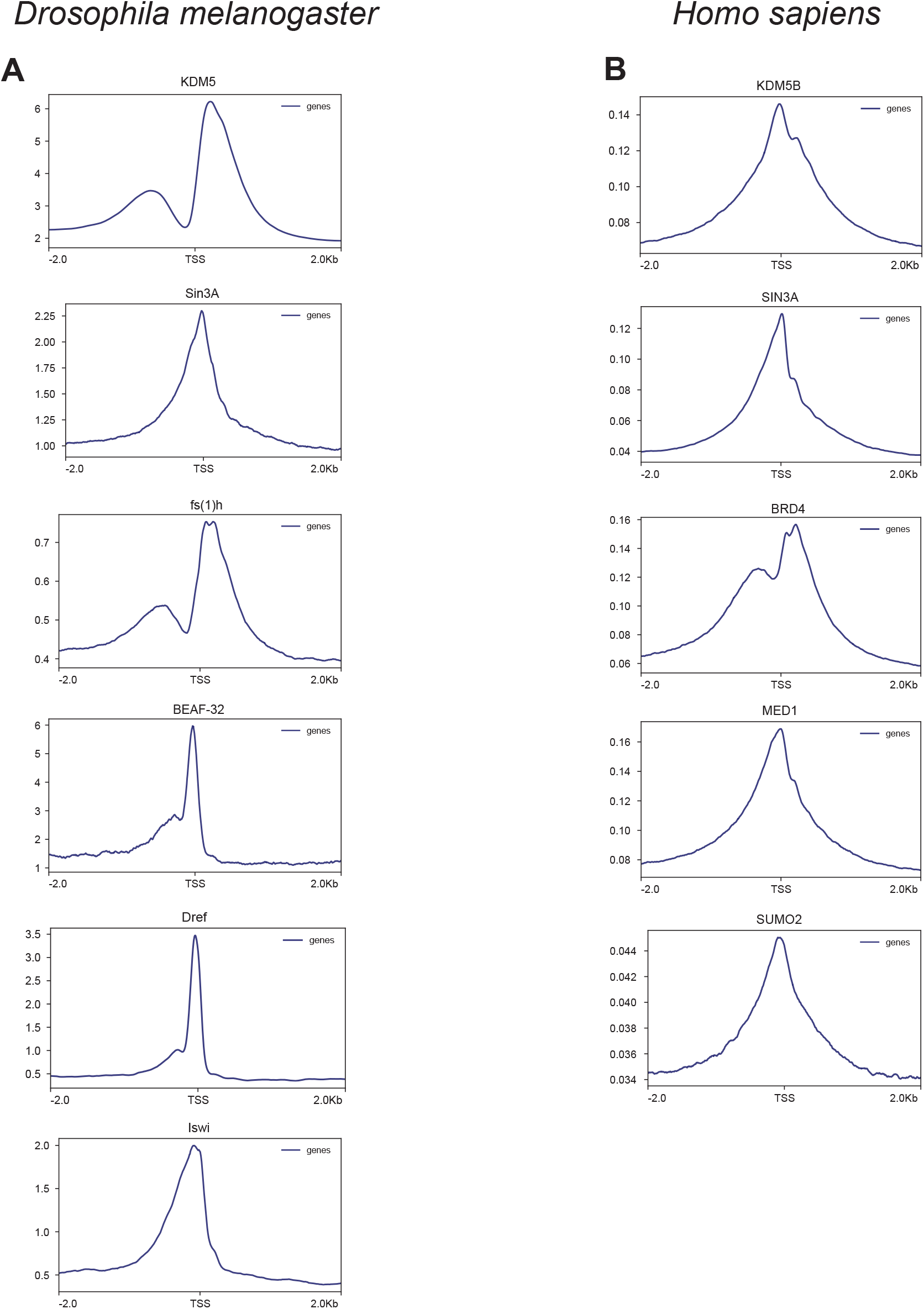
Binding of KDM5 proteins and identified interactors relative to the transcriptional start site. (A) Genomic binding profiles of *Drosophila* KDM5, Sin3A, fs(1)h, BEAF-32, Dref and Iswi relative to the TSS. (B) Genomic binding profiles of human KDM5B, SIN3A, BRD4 (fs(1)h ortholog), MED1 and SUMO2 relative to the TSS.

## MATERIALS AND METHODS

### Fly strains and care

Fly crosses were maintained at 25°C with 50% humidity and a 12-hour light/dark cycle. Food (per liter) contained 18g yeast, 22g molasses, 80g malt extract, 9g agar, 65 cornmeal, 2.3g methyl para-benzoic acid, 6.35ml propionic acid. The number of male and female animals were equal across all genotypes examined. The *kdm5*^*140*^ null allele has been previously described. (33)

### Cloning and Transgenesis

The N- and C-terminally TurboID-tagged constructs were generated by cloning the coding region of *kdm5* upstream or downstream of HA:TurboID from pCDNA3-TurboID (RRID:Addgene_107171)(46) in the pUASpattB vector (RRID:DGRC_1358). UASp-HA:TurboID with a NLS was generated by cloning HA:TurboID into the same UASpattB vector. UASp-dCas:HA:Turbo:NLS was made by combining the dCas9 open reading frame from SID3s-dCas9-KRAB (RRID:Addgene_106399) with HA:TurboID:NLS in the pUASpattB vector (RRID:DGRC_1358). All transgenes were generated by injection into y^1^ M{RFP[3xP3.PB] GFP[E.3xP3]=vas-int.Dm}ZH-2A w^*^; M{3xP3-RFP.attP}ZH-86Fb at BestGene Inc.

### Western Blotting

Western analyses were carried out as previously described. (33) Briefly, five 2- to 5-day old adult fly heads were homogenized in 2x NuPAGE LDS sample buffer, sonicated for 10 mins, treated with DTT, run on a 4-12% Bis-Tris 1 mm gel and transferred to a PVDF membrane. The following primary antibodies were used: mouse anti-HA (1:1000, Cell Signaling Technology Cat# 2367, RRID: AB_10691311), Streptavidin 680 (1:10,000, ThermoFisher, Streptavidin Alexa Fluor 680 conjugate), rabbit anti-alpha-Tubulin (1:5000, Cell Signaling Technology Cat# 2144, RRID:AB_2210548). Secondary antibodies used were IRDye^®^ 680RD Donkey anti-Mouse IgG (1:5000; LI-COR Biosciences Cat# 925-68072, RRID: AB_2814912) and IRDye^®^ 800CW Donkey anti-Rabbit IgG (1:5000; LI-COR Biosciences Cat# 926-32213, RRID: AB_621848). Blots were scanned and processed using a LI-COR Odyssey Infrared scanner.

### Purifying and identifying proteins using TurboID Biotinylated Protein Enrichment

2-5 day-old flies were flash frozen in liquid nitrogen and decapitated and a total of ten heads were used per sample. Heads were homogenized in 250 µL RIPA Buffer (Thermofisher 89901) supplemented with Halt™ Protease Inhibitor Cocktail (Thermofisher, 78430) and centrifuged at 4 °C for 10 minutes at 15,000XG to remove debris. 100 µL of Pierce™ Streptavidin Magnetic Beads (Thermofisher, 88817) were washed twice with RIPA and the cleared lysate was added. The lysate-bead mixture was incubated with rotation at 4 °C overnight. The next day the lysate was discarded, and beads were washed twice with RIPA, once with 1M KCl, once with 0.1 M Na_2_HCO_3_, once with 1 M Urea in 10 mM Tris pH 8.0, and twice again with RIPA. For Western Blot analyses all RIPA was removed and biotinylated proteins were eluted with 4X NuPAGE™ LDS Sample Buffer (Invitrogen, NP0007) supplemented with 2 mM biotin and 20 mM DTT.

### On-bead protein digestion

Proteins were digested directly on streptavidin beads. 5 mM DTT and 50 mM ammonium bicarbonate (pH = 8) were added to the solution and left on the bench for about 1 hour for disulfide bond reduction. Samples were then alkylated with 20 mM iodoacetamide in the dark for 30 minutes. Afterward, 500 ng of trypsin was added to the samples, which were digested at 37 °C for 18 h. The peptide solution was dried in a vacuum centrifuge.

### Sample desalting

Prior to mass spectrometry analysis, samples were desalted using a 96-well plate filter (Orochem) packed with 1 mg of Oasis HLB C-18 resin (Waters). Briefly, the samples were resuspended in 100 µl of 0.1% TFA and loaded onto the HLB resin, which was previously equilibrated using 100 µl of the same buffer. After washing with 100 µl of 0.1% TFA, the samples were eluted with a buffer containing 70 µl of 60% acetonitrile and 0.1% TFA and then dried in a vacuum centrifuge.

### LC-MS/MS Acquisition and Analysis

Samples were resuspended in 10 µl of 0.1% TFA and loaded onto a Dionex RSLC Ultimate 300 (Thermo Scientific), coupled online with an Orbitrap Fusion Lumos (Thermo Scientific). Chromatographic separation was performed with a two-column system, consisting of a C-18 trap cartridge (300 µm ID, 5 mm length) and a picofrit analytical column (75 µm ID, 25 cm length) packed in-house with reversed-phase Repro-Sil Pur C18-AQ 3 µm resin. Peptides were separated using a 90 min gradient from 4-30% buffer B (buffer A: 0.1% formic acid, buffer B: 80% acetonitrile + 0.1% formic acid) at a flow rate of 300 nL/min. The mass spectrometer was set to acquire spectra in a data-dependent acquisition (DDA) mode. Briefly, the full MS scan was set to 300-1200 m/z in the orbitrap with a resolution of 120,000 (at 200 m/z) and an AGC target of 5×10e5. MS/MS was performed in the ion trap using the top speed mode (2 secs), an AGC target of 1×10e4 and an HCD collision energy of 35. Raw files were searched using Proteome Discoverer software (v2.4, Thermo Scientific) using SEQUEST search engine and the UniProt database of Drosophila melanogaster. The search for total proteome included variable modification of N-terminal acetylation, and fixed modification of carbamidomethyl cysteine. Trypsin was specified as the digestive enzyme with up to 2 missed cleavages allowed. Mass tolerance was set to 10 pm for precursor ions and 0.2 Da for product ions. Peptide and protein false discovery rate was set to 1%. Following the search, data was processed as described by Aguilan et al.(98). Briefly, proteins were log2 transformed, normalized by the average value of each sample and missing values were imputed using a normal distribution 2 standard deviations lower than the mean. Statistical regulation was assessed using heteroscedastic T-test (if *p*-value < 0.05). Data were assumed to be Gaussian distributed but this was not formally tested.

### Interaction Map Generation

STRINGDB(62) and Cytoscape(63) were used for physical interaction mapping. Lines between proteins represent physical interaction and the darkness of the lines represent the confidence of physical interaction. A confidence score of greater than 0.4/1 was used as a cutoff. Nodes were manually positioned and annotated using STRING GO Clusters and published literature as an organizational guide.

### Bioinformatic Analyses

Gene Ontology analysis utilized R packages clusterProfiler (v4.4.4)(99) and ReactomePA (v1.40.0)(100). Volcano plots were generated using EnhancedVolcano (v1.14.0).(101) The Enrichment Analysis function on ChIP Atlas(74) was used to perform permutation tests, which compares the overlap of datasets using genomic ranges of called peaks (BED files). For our studies, the query datasets were *Drosophila* KDM5 (SRX1084165) and Human KDM5B (SRX3285561), which were compared using the following specific parameters: “TFs and others”, cell type class was set to ‘All types’ and ‘Breast’ for *Drosophila* and Human respectively. A 100X random permutation of each was used as the control. For these analyses, a single base pair overlap is considered as an overlap. For *Drosophila*, selected profiles were generated using bigWig files from: KDM5 (SRX1084165), Sin3A (SRX1158165), BEAF-32 (SRX386677), Dref (SRX749042), Iswi (SRX5346167), and fs(1)h (SRX203000). For Human, selected profiles generated using: KDM5B (SRX3285561), SIN3A (SRX190318), MED1 (SRX673749), BRD4 (SRX5089551), and SUMO2 (SRX3541112). Deeptools(3.5.1)(102) computeMatrix and plotHeatmap functions were used to make profiles and heatmaps. For these, the corresponding BED files from each interactor’s SRX accession was used as the --region option. The bigWigs for the interactor and KDM5 were used in the –score option, bin size was set to 5 bp. Due to large differences in peak number, for Figure 4K, KDM5B’s BED file was used as the region file. Pygenometracks(3.7)(103) was used to visualize ChIP-seq tracks.

